# Solitary bee larvae prioritize carbohydrate over protein in parentally provided pollen

**DOI:** 10.1101/397802

**Authors:** Alexander J. Austin, James D. J. Gilbert

## Abstract

1. Most organisms must regulate their nutritional intake in an environment full of complex food choices. While this process is well understood for self-sufficient organisms, dependent offspring, such as bee larvae, in practice have limited food choices because food is provided by parents. Nutrient balancing may therefore be achieved by offspring, by parents on offspring’s behalf, or by both, whether cooperatively or in conflict.
2. We used the Geometric Framework to investigate the capacity of dependent larval mason bees (*Osmia bicornis*) to regulate their intake of protein and carbohydrate. Female *Osmia* seal eggs individually inside cells they have provisioned with pollen, and have no contact with developing offspring, allowing offspring choices to be studied in isolation. Herbivorous insect larvae are typically expected to balance protein and carbohydrate to maximise growth and reproduction.
3. Contrary to prediction, carbohydrate and not protein mediated both growth and survival to pupation. Accordingly, larvae prioritised maintaining a constant intake of carbohydrate and self-selected a relatively carbohydrate biased diet compared to other hymenopterans, while tolerating wide excesses and deficiencies of protein, rendering them potentially vulnerable to dietary change or manipulation. Reasons for prioritising carbohydrate may include (1) the relative abundance of protein in their normal pollen diet, (2) the relative paucity of nectar in parental provisions making carbohydrate a scarce resource, or (3) the requirement for diapause for all *O. bicornis* larvae. Larvae were intolerant of moderate dietary dilution, likely reflecting an evolutionary history of nutrient-dense food.
4. Our results demonstrate that dependent offspring can remain active participants in balancing their own nutrients even when sedentary, and, moreover, even in mass provisioning systems where parents and offspring have no physical contact. Research should now focus on whether and how evolutionary interests of parent and dependent offspring coincide or conflict with respect to food composition, and the implications for species’ resilience to changing environments.

## Introduction

Most animals manage their nutrient intake by combining nutritionally different foods (Simpson & Raubenheimer 2012). However, the importance of this ability depends upon the nutritional variability of the animals’ typical food (Despland & Noseworthy 2006; Raubenheimer, Simpson & Mayntz 2009). Extreme specialists, for example, can lose the capacity to regulate nutrition (Warbrick-Smith *et al*. 2009; Poissonnier *et al*. 2018). One way in which organisms can experience limited nutritional choice is if they are dependent upon others for nutrition, or “alloregulation” (Lihoreau *et al*. 2014), such as dependent offspring of altricial birds, human toddlers, and many larval insects. Under these circumstances, by what rules offspring regulate their own consumption should depend upon provisioning rules of parents. On the one hand, offspring tend to have different requirements from parents (Harper & Turner 2000; Michaelsen *et al*. 2003) - particularly for protein, given their elevated rates of somatic growth and development. Accordingly, parents often make different nutritional choices for their offspring versus when foraging for themselves (Royama 1970; Dussutour & Simpson 2009; Burt & Amin 2014). For example, granivorous birds usually provision young with insects, rather than the seed diets of adults, to fulfil protein requirements (Wiens & Johnston 2012). If parents alloregulate offspring nutrition tightly, then offspring should have no need for self regulation, like extreme specialists (Poissonnier *et al*. 2018). On the other hand, parents may provide suboptimal nutrition for offspring - either through inefficiency (e.g. Seidelmann 2006), or if parents’ and offspring’s evolutionary interests do not coincide (Trivers 1974). Here, offspring may be able to use nutritional regulation to mitigate costs arising from their parents’ nutritional choices. While there has been much research into evolutionary compromises involving offspring solicitation and corresponding parental responses (e.g. Smiseth, Wright & Kölliker 2008), less is known about whether or how offspring may exert control by discriminating among parental provisions.

The Geometric Framework for Nutrition (GF) allows us to investigate foraging decisions made by animals in multi-dimensional “nutrient space” (Simpson & Raubenheimer 1993). The GF can be used to determine animals’ nutritional choices relative to their “intake target” - the optimal amount and balance of multiple macronutrients - as well as their “rule of compromise” that governs their choices when restricted to suboptimal food (Raubenheimer & Simpson 1999b). The GF has provided insights into the nutritional ecology of a broad range of taxa (reviewed in Simpson & Raubenheimer 2012). Its application to dependent offspring, though, has typically been as part of studies of social insect systems (e.g. Helm *et al*. 2017) and studies have often inferred offspring requirements indirectly from patterns of alloparental feeding in studies more broadly focused on adult foraging (see Dussutour & Simpson 2009; Cook *et al*. 2010; Vaudo *et al*. 2016). In such systems, multiple adults normally contact offspring, progressively feeding and adjusting nutrition in response to feedback (Field 2005; Schmickl & Karsai 2017), making the responses of individual larvae difficult both to follow and interpret.

In solitary bees, by contrast, typically females provision offspring individually with a pollen ball before sealing the cell and leaving. This behaviour makes solitary bees an ideal, manipulable model for directly studying the nutrition of dependent larvae (Strohm *et al*. 2002) independently of provisioning decisions made by parents. Larvae of bees, like most aculeate hymenopterans, rely on parents or alloparents for nutrition (Field 2005). Nutritional requirements for bee adults and offspring differ, often radically (Weeks *et al*. 2004; Filipiak 2019); adults primarily feed on carbohydrate-rich nectar (although see Cane 2016) while larvae feed mostly on protein-rich pollen (Filipiak 2019). Solitary bees, along with most other hymenopterans and many other parental insects, typically have a simple one-to-one parent-offspring relationship whereby parents “mass provision” their young, providing a finite, fixed-mass food provision, and have no contact with their young during development (Costa 2006). Such systems are almost unstudied in a rigorous nutritional context (but see e.g. Roulston & Cane 2002). In these species, there is no opportunity for parents to adjust nutrition according to offspring feedback, and the larva must therefore make the best of what it is given. It may be that offspring regulate their own nutrition to compensate for variation, as in more independent insect larvae (Lee *et al*. 2002), or possibly to mitigate costs imposed by parents. Alternatively, they may have lost this capacity, like extreme specialists (Warbrick-Smith *et al*. 2009; Poissonnier *et al*. 2018). We know very little about how bee larvae deal with variable nutrition (but see Helm *et al*. 2017) - a knowledge gap with potentially important consequences, considering the proposed link between nutritional stress and bee declines (Roulston & Goodell 2011; Goulson *et al*. 2015).

In this study, we used a commercially important solitary bee species, *Osmia bicornis*, to investigate how dependent larvae cope with varying nutrition, and whether they can regulate their own intake. *O. bicornis* are pollen generalists (Falk 2015) and the solid, roughly spherical pollen balls that parents provide to offspring are variable in species composition (Haider *et al*. 2014). Although *O.bicornis* larvae are sedentary, they are capable of limited movement, in principle allowing them to preferentially consume specific parts of a fixed provision (note that other invertebrates are also capable of extracting and consuming preferred nutrients from nutritionally complex food items (Mayntz *et al*. 2005). The species is entirely solitary with no known tendency for offspring to “help at the nest” as in other bees (Hogendoorn & Velthuis 1993; Rehan *et al*. 2014) so there is no reason to believe mothers would alter offspring nutrition to force them to help, as in other systems (Lawson, Helmreich & Rehan 2017) and therefore no obvious potential for parent-offspring conflict over offspring nutrition. Natural variation in pollen ball nutrient content is largely unquantified (although see Budde & Lunau 2007), so there is no prior expectation about the capacity of larvae to regulate their consumption. We used a classic GF design (e.g. Lee *et al*. 2008), focusing on protein and carbohydrate, with two experimental phases. In the first “no-choice” phase we raised *O. bicornis* larvae on fixed diets of differing protein to carbohydrate ratios to determine their rules of compromise and the diet composition that maximised fitness. In a second “choice” phase, we then provided larvae with targeted choices between sets of two imbalanced diets that differed in their protein:carbohydrate ratios to determine whether larvae defend an intake target. Given the central role of protein in growth of insect larvae (Scriber & Slansky 1981; Behmer 2009), and following Hunt & Nalepa’s (1994) exhortation to “follow the protein”, we predicted that (1) protein would be a key driver of fitness in larval *O. bicornis*, (2) larvae would accordingly aim for a relatively protein-biased intake target, and (3) larvae would prioritize regulating intake of protein over carbohydrate.

## Methods

### Study organism

*Osmia bicornis* is a common, cavity-nesting solitary bee (Falk 2015), and a commercially important pollinator (Jauker *et al*. 2012). *O. bicornis* larvae were obtained as diapausing adults in cocoons (Mauerbienen®). These were released at the nesting site at the University of Hull in April 2017, and emerging adults allowed to breed. Early trials revealed that fresh eggs and newly emerged larvae were too fragile for manipulation. Therefore, newly emerged larvae were left alone for two days before we transferred them to a single-occupancy nest and assigned each to an experimental treatment. Details of nesting apparatus and monitoring protocols are available in the supplementary methods.

### Diet Formulation

Existing artificial diet protocols for solitary bees have met with limited success in terms of larval survival (Nelson, Roberts & Stephen 1972; Fichter, Stephen & Vandenberg 1981). We used six diets, consisting of three different protein:carbohydrate (P:C) ratios (Diet A = 1:1.2, Diet B = 1:2.3 & Diet C = 1:3.4) and two total macronutrient concentrations (concentration 1 = 90% nutrients, 10% diluent, or concentration 2 = 70% nutrients, 30% diluent; see table S1). Diet ratios were chosen based on a combination of the nutrient ratios in honeybee-collected pollen loads and published data for protein content of *O. bicornis* pollen balls (Budde & Lunau 2007). Diets were diluted with sporopollenin, the primary constituent of the exine of pollen (Mackenzie *et al*. 2015), an extraordinarily stable natural polymer. Sporopollenin is a novel dietary diluent for bees; its suitability has been demonstrated in a separate study (Tainsh *et al*. 2020). For a more detailed description of sporopollenin and its preparation, see supplementary methods.

### Experiment 1: No-choice phase

Two-day-old larvae, randomized by parentage, were allocated to one of 6 treatments corresponding to our 6 artificial diets (n = 20/treatment). Provisions were made to resemble the size of natural provisions (mean initial artificial provision weight = 0.323g +/- 0.034g). Once provisioned, larvae were placed in an incubation chamber (Gallenkamp, IH-270) at 23°C and 80% RH. Provisions were replaced weekly to avoid desiccation and mould formation, or when fully consumed by larvae, ensuring the diet was always available in excess. Weight of provision consumed was recorded upon provision replacement. A “water control” group, containing artificial diets but no larvae, was used to track water loss from the diets, going through the same weighing regime as above with weight loss recorded at each swap. Nests were checked daily for mortality. Cocoon weight was recorded at the completion of pupation.

### Experiment 2: Choice phase

In the choice experiment, 32 two-day-old larvae of mixed parentage were randomly divided among four treatments. Treatments consisted of strategic pairwise combinations (see Fig 1; Table 1) of four possible diets: A1 (1P:1.2C, 90%), A2 (1P:1.2C, 70%), C1 (1P:3.4C, 90%), C2 (1P:3.4C, 70%). Because *O. bicornis* larvae are sedentary and receive a single provision, it is not biologically appropriate to present choices between two diets simultaneously. Therefore, choices were offered temporally by swapping the provision every other day, presenting one diet at a time. This required the larvae to differentially feed over time to compensate for temporal imbalance, in order to converge on an intake target (see e.g. Raubenheimer & Jones 2006). All larvae were kept on the same treatment from two days post-hatching up to pupation, whereupon diet replenishment ceased. The diet that the larvae would be fed first was randomly assigned via coin toss prior to the experiment.

**Figure 1.**
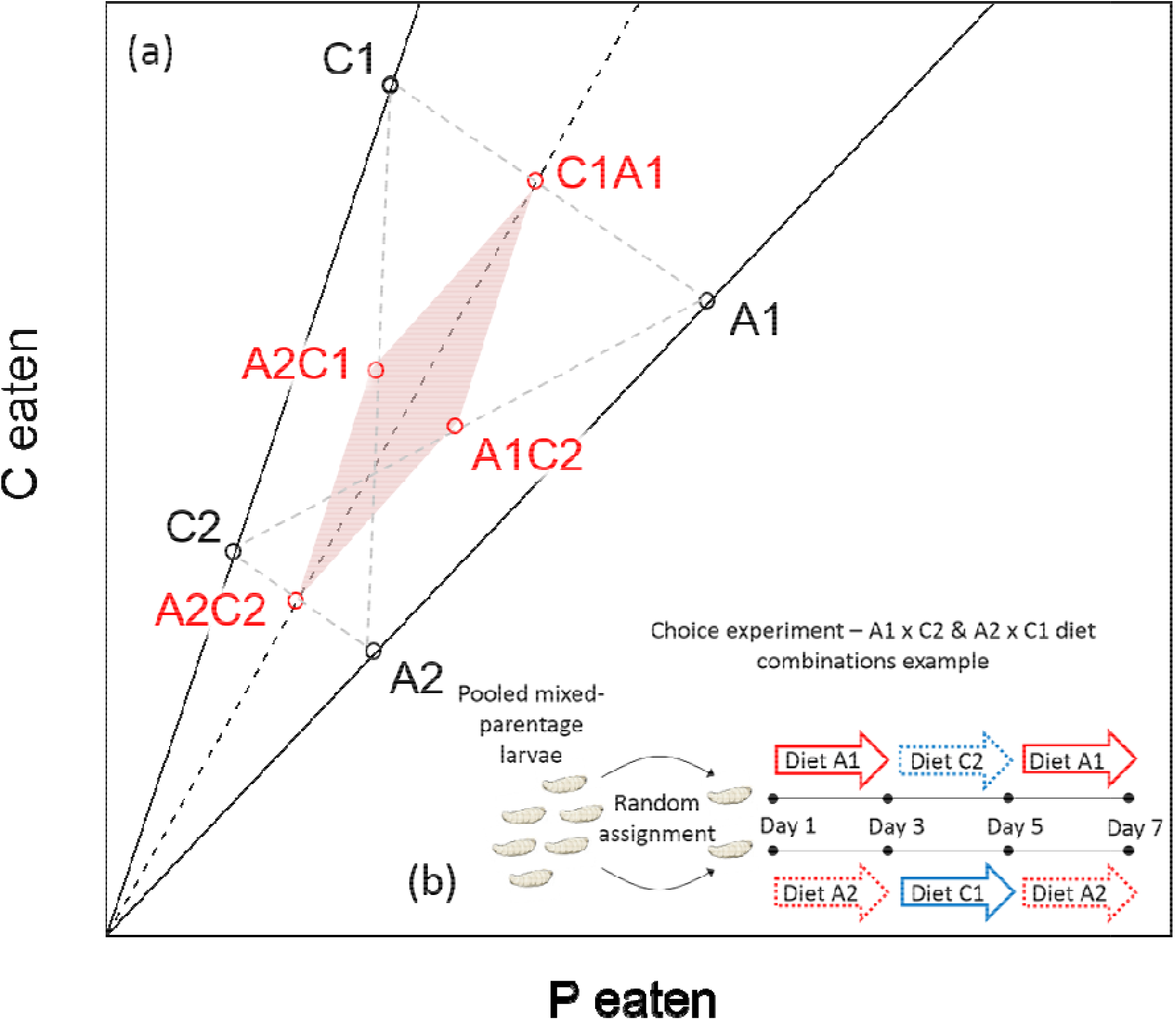
**(a)** Expected protein and carbohydrate consumption if larvae ate indiscriminately between two diets. Diet choices are pairwise combinations of diets A1, A2, C1 and C2, which each contain protein and carbohydrate at different ratios and concentrations. Solid lines represent P:C ratios; black points represent actual nutrient content of each diet, which depends upon dilution as well as P:C ratio. Red points represent expected consumption if larvae eat randomly (i.e. equally) from each of a choice of two diets (choices denoted by the red point labels). **(b)** schematic describing how larvae were assigned to each diet grouping. Coloured arrows show the period in days that each larva was fed a particular diet.

**Table 1.** Sample sizes for each diet combination used for choice phase (allocated by random coin toss). “Order” refers to diet order - e.g. for A1C1, Order 1 would receive A1 first whereas Order 2 would receive C1 first, determined by coin toss. Surviving larvae are in parentheses.

|  | Order 1 | Order 2 | Total |
| --- | --- | --- | --- |
| A1C1 | 1 (1) | 6 (5) | 7 (6) |
| A1C2 | 5 (5) | 4 (3) | 9 (8) |
| A2C1 | 3 (2) | 5 (5) | 8 (7) |
| A2C2 | 5 (2) | 3 (1) | 8 (3) |

### Statistical Analysis

All analyses were conducted in R version 3.4.2 (R Core Team, 2017). For the no-choice experiment, we calculated total nutrients consumed (protein and carbohydrate) from raw diet consumption data for each swap, adjusted for water loss and dilution. Values were then summed for each larva.

To investigate consumption rules, including rules of compromise, we first asked whether diet ratio and concentration affected consumption of (a) the total provision, (b) protein, or (c) carbohydrate, using models of each respective variable with “ratio” and “concentration” as predictors. Rules of compromise can include nonlinear effects, particularly curves around the intake target (Simpson & Raubenheimer 1993). To account for potentially curvilinear relationships we also added ratio^2^ as a predictor, as well as two-way interactions between all predictors.

To assess fitness consequences of macronutrient consumption, we analysed cocoon weight at pupation and survival to pupation. For both analyses, to analyse potentially nonlinear effects of nutrient consumption upon fitness, we used polynomial regression, fitting both first- and second-order polynomial terms for “protein consumed [P]” and “carbohydrate consumed [C]”. We analysed cocoon weight using a linear model with “cocoon weight” as a response. The full model contained linear (P and C) and quadratic effects for both nutrients (P^2^ and C^2^) and their interaction (P × C), as well as diet concentration (high or low), and two-way interactions between concentration and nutrients (conc × P, conc × P^2^, conc × C, conc × C^2^). We used standard diagnostics to check the fit of models, and used a reverse stepwise process to determine the minimal model, at each step dropping the least significant term until the model contained only significant terms. To analyse survival, we used parametric survival analysis in the *survival* package in R and fitted the same full model as described above. We assessed model fit graphically by inspecting the Kaplan-Meier estimates of the residuals against the assumed Weibull distribution. Again we used reverse stepwise selection to determine the minimal model, comparing models with likelihood ratio tests against a chi squared distribution. To visualise these fitness effects, we calculated response surfaces for cocoon weight and survival, and visualised them using non-parametric thin-plate splines.

In the choice experiment, the mean final consumption of each nutrient was investigated using linear models with diet combination, dilution and their interaction as predictors, and using Tukey’s post hoc tests to compare individual treatments against each other. Under a null expectation we would expect larvae to eat randomly from each diet (Fig 1). Thus, for each larva we calculated the deviation from this null expectation. We then tested whether these values systematically departed from zero for protein and carbohydrate, and whether these departures from random consumption differed by treatment group. We used a linear model with “deviation from random consumption” as the response variable and “treatment group” as a predictor.

Larvae that died pre-pupation were not used in the calculation of the mean protein and carbohydrate consumption for diets in either experiment, or for cocoon weight, but were used in analyses involving survival.

## Results

### No-choice phase

Dietary P:C ratio had a significant effect on the overall amount of provision consumed, with larvae consuming more provision on high P:C ratio diets (F_1,78_=21.55, p<0.0001). Total consumption was also affected by diet concentration (F_1,78_=14.03, p<0.001); larvae on less concentrated diets consumed more provision, indicating compensatory feeding. The quadratic term was not significant (Fig. 2a; Table S2a).

Dietary P:C ratio had a strong effect on the total amount of P eaten (F_1,79_=146.93, p<0.0001); more protein was eaten by larvae raised on the higher P:C diets (Fig. 3). Diet concentration had no effect on the amount of P consumed; neither was there a ratio:concentration interaction, nor a quadratic effect of ratio (Table S2b).

**Figure 2.**
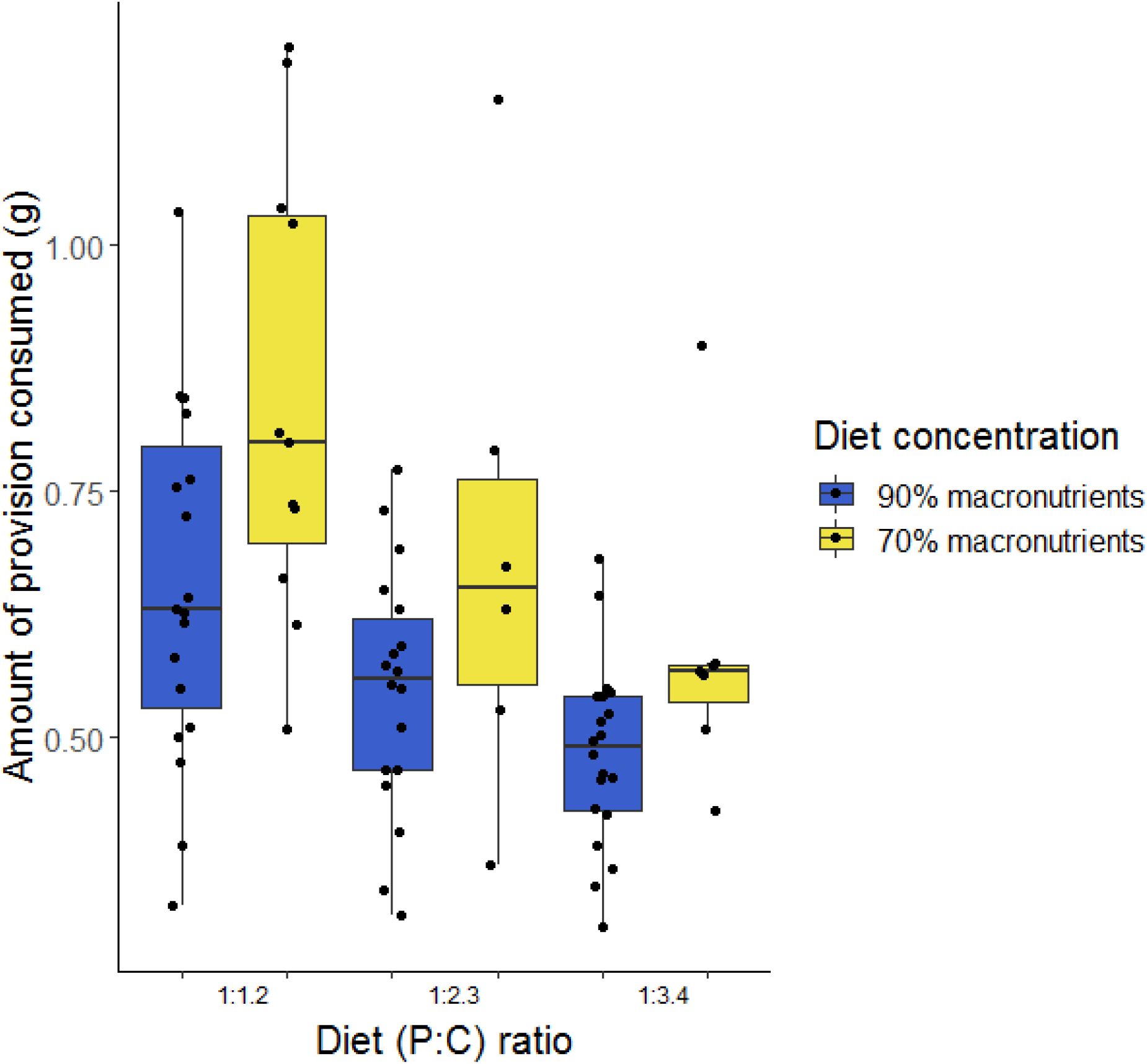
Amount of provision in grams consumed by larvae raised on the 3 different P:C ratio artificial diets at the 2 different macronutrient concentrations (90% and 70% macronutrient content).

**Figure 3.**
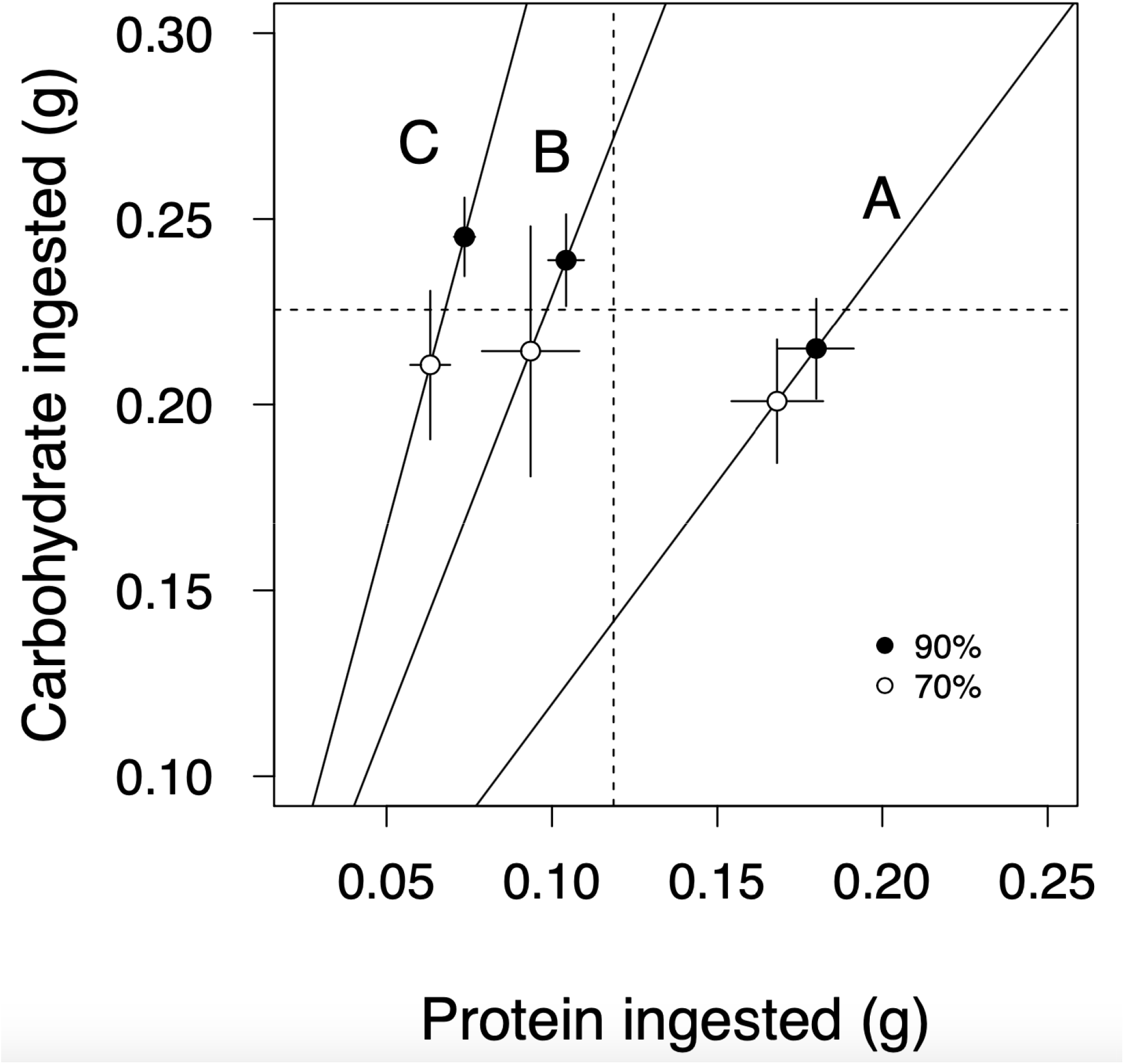
Mean total (+/-1 SE) amount of P and C consumed in grams by larvae on each diet before pupation. Solid lines and letters represent three P:C ratios (A = 1:1.2, B = 1:2.3, C = 1:3.4). Numbers following letters denote diet concentration (1 = 90%, 2 = 70%). Dotted lines show global mean consumption of each nutrient.

In contrast, larvae consumed similar amounts of C across all diets, with neither concentration nor dietary P:C ratio (linear or quadratic) having an influence on the amount of C consumed (Table S2c). A mean of 0.23 ± 0.01 g of C was consumed by (surviving) larvae across all diet treatments (Fig. 3).

Cocoon weight varied differently with macronutrient intake depending on the overall concentration of the diet (carbohydrate × conc interaction, F_1,72_=6.50, p=0.01; protein × conc interaction, F_1,72_=4.82, p=0.03). At 90% nutrient density, cocoon weight was correlated positively with the amount of carbohydrate consumed, and negatively with protein (Fig. 4a). For our range of diets, the greatest weights were obtained by larvae that ate above approx. 0.3g C and below 0.15g P. In contrast, at 70% nutrient density, cocoons were lower in weight than on the 90% diets, and were fairly uniform in weight irrespective of macronutrient intake (Fig 4b). No quadratic effects were observed, nor interactions involving quadratic effects, meaning that we did not identify an optimal amount of P or C that maximised cocoon weight within the range of diets we used (Table S3a).

**Figure 4.**
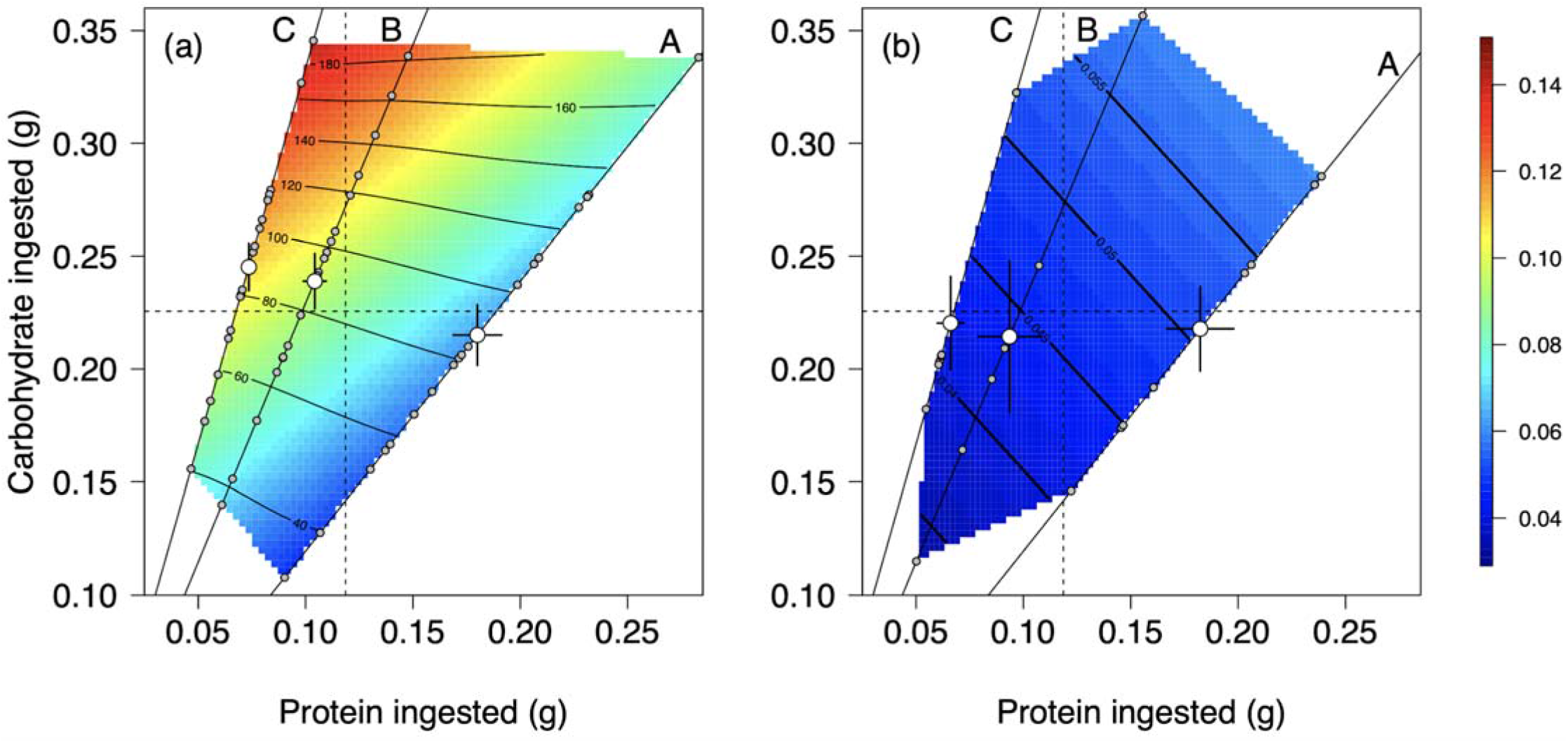
Effects of P and C consumption upon cocoon weight (g) in larvae fed diets at (a) 90% and (b) 70% nutrient density. Transition from blue to red indicates heavier cocoons. For context, mean total consumption of P and C for each diet is plotted (white points; data as in Fig. 2) alongside raw data (grey points). Solid lines and letters represent three P:C ratios (A = 1:1.2, B = 1:2.3, C = 1:3.4).

The relationship between survival and nutrition similarly depended upon dietary concentration (carbohydrate × conc interaction: χ_1_=6.50, p=0.01). Survival of larvae fed diets at 90% concentration depended primarily upon carbohydrate consumption (Fig 5a). Those larvae that consumed high amounts of carbohydrate saw the highest survival irrespective of how much protein was consumed. At lower levels of carbohydrate, interestingly, protein weakly mediated survival (protein × carbohydrate interaction: χ_1_=−4.88, p=0.046). Survival of larvae raised on the more dilute diets was much lower, and was not substantially affected by intake of P or C (Fig 5b). Again, there were no significant quadratic terms, whether as main effects or as part of interactions (Table S3b).

**Figure 5.**
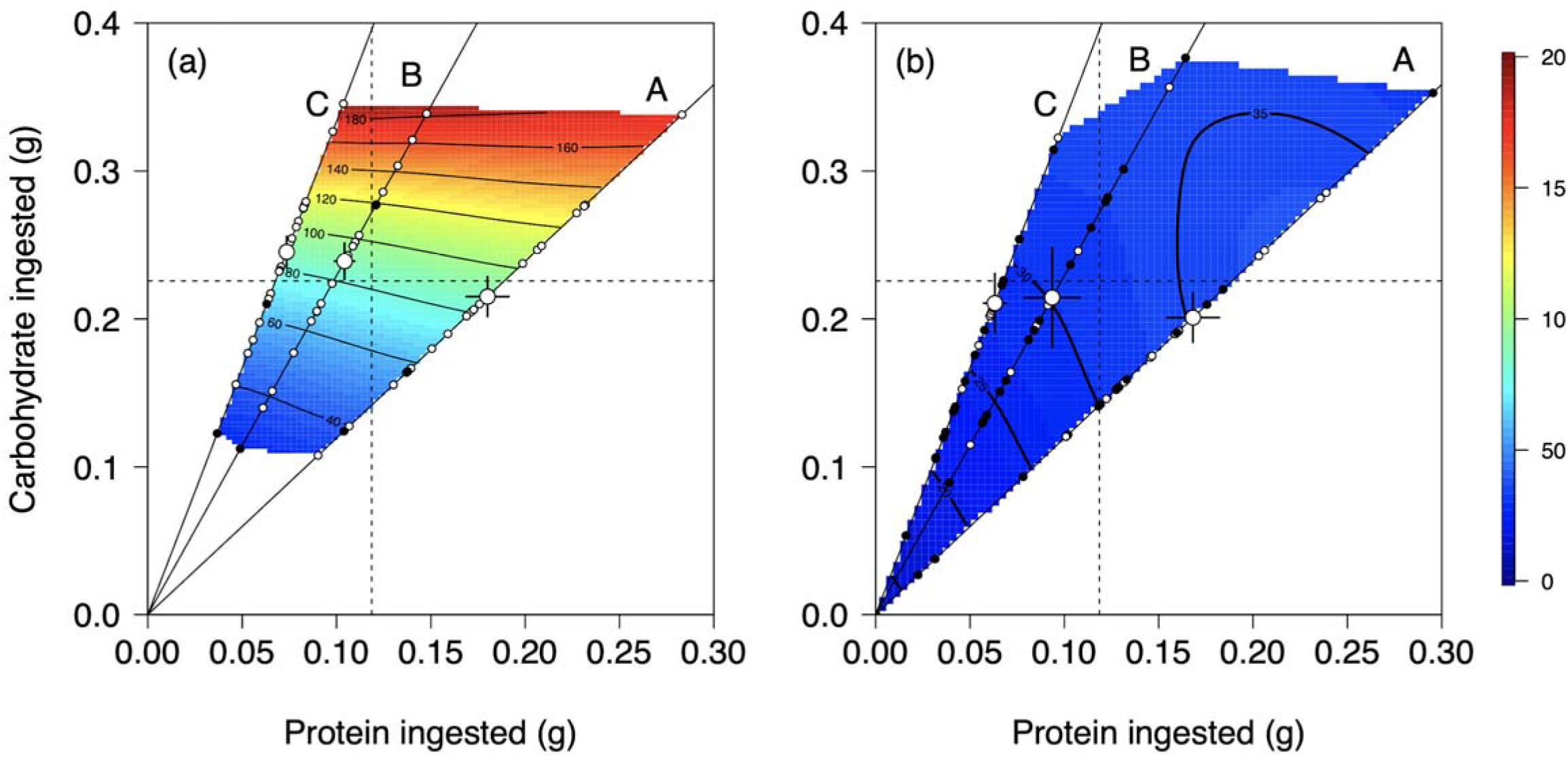
Effects of P and C consumption upon estimated survival time (colour) in larvae fed diets at (a) 90% and (b) 70% nutrient density. Transition from blue to red indicates longer survival. Black points, dead larvae; white points, larvae surviving to pupation. For context, mean total consumption of P and C for each diet is plotted (large white points; data as in Fig. 2). Solid lines and letters represent three P:C ratios (A = 1:1.2, B = 1:2.3, C = 1:3.4).

### Choice phase

We found no evidence of larvae defending a common intake target *sensu stricto (Raubenheimer & Simpson 1993, 1999b*), i.e. a common ratio *and* amount of nutrients consumed, which would have been evident as all groups clustering at a common point in nutrient space in Fig 6a. Nevertheless, consumption deviated from random so as to converge upon a target P:C ratio (see e.g. Deans, Sword & Behmer 2019) represented by a common line, or “nutritional rail”, of approx. 1:1.8 (Fig. 6a). The amount of protein consumed by larvae was significantly affected by diet combination: more protein was consumed by individuals offered diet combinations that were overall more concentrated (F_3,23_=7.43, p<0.01, Fig 6a; Table S4a). Similarly, carbohydrate consumption was significantly affected by diet combination (F_3,23_=4.58, p=0.01, Fig 6a; Table S4b). Unlike with protein, though, this pattern appeared to be driven by the diets at the extreme; only the most concentrated diet pair (C1A1) differed from the least concentrated pair (C2A2; Fig 6a); other pairwise comparisons were not significant (Table S4).

**Figure 6.**
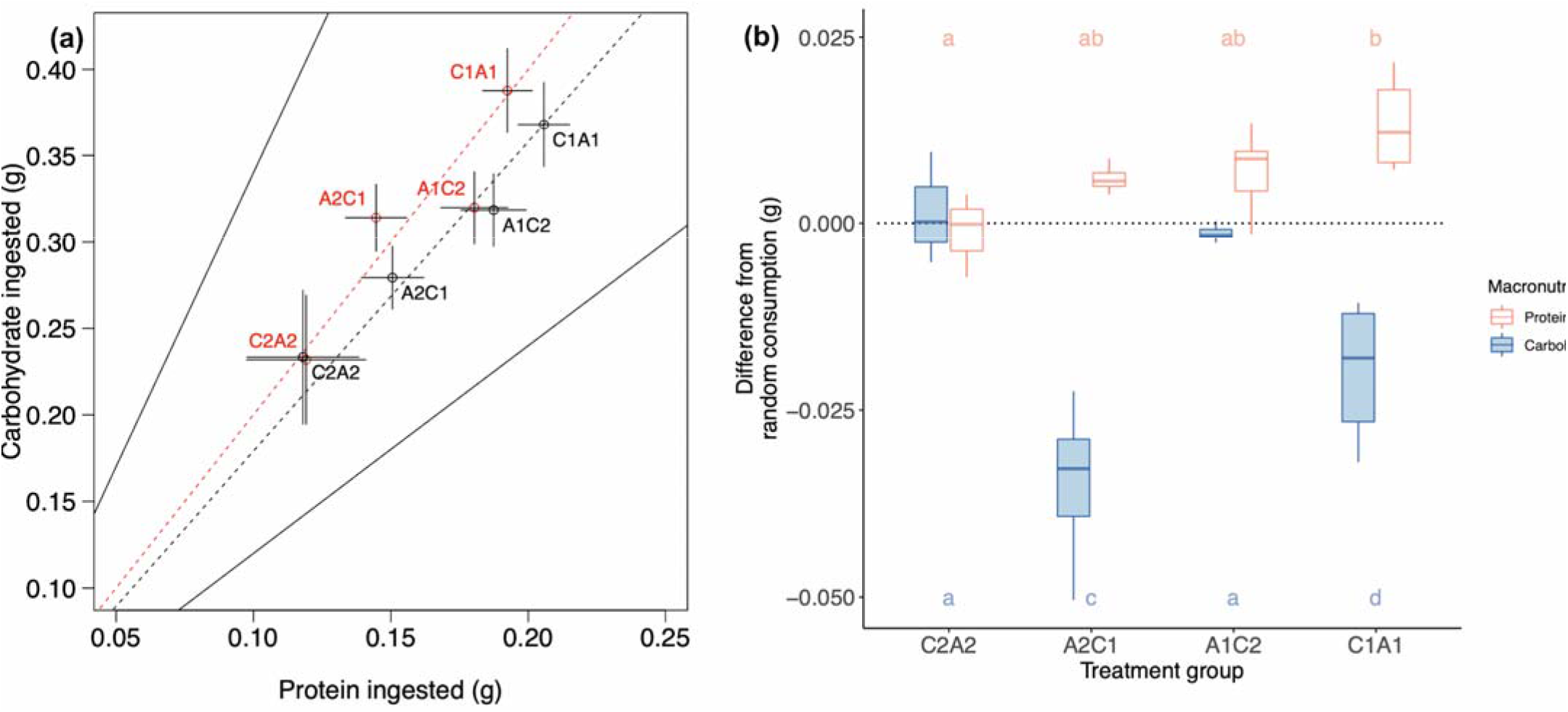
**(a)** The mean (+/- SE) amount of protein (P) and carbohydrate (C) eaten by larvae in the choice experiment. Each point label denotes a choice of two diets, one A and one C; black labels show observed intake, red labels show expected intake under random consumption. Letters represent diet P:C ratio (A = 1:1.2, C = 1:3.4); numbers represent diet concentration (1 = 90%, 2 = 70%), hence, for example, “A2C1” represents the pairing of diet A2 with diet C1. Solid lines represent dietary P:C ratios (Top line = Diet C, Bottom line = Diet A). Dashed red line shows expected average P:C ratio based on random consumption. Dashed black line shows average P:C ratio of observed intake across larvae, (b) Deviation from random intake of protein and carbohydrate for larvae in different treatment groups during the choice phase. Treatment groups are given in order of overall diet concentration. Bars with similar letters displayed above or below are not statistically significantly different (Tukey’s post-hoc comparisons).

Despite the lack of a common intake target, larvae were not consuming diets at random (Fig 6b, Table S4c, d). For both carbohydrate and protein we saw differences in consumption from what would have been expected for each larva based on random consumption, and this effect was dependent on the specific set of diet choices (protein, F_4,20_=19.67, p<0.001; carbohydrate, F_4,20_=51.65, p<0.001). When visualised as the amounts of protein and carbohydrate consumed during each 48h treatment period (Fig. 7), it is clear that larvae were achieving a degree of homeostasis in carbohydrate consumption (Fig 7b) compared to what would be expected under random consumption of each diet choice (Fig 7a), whereas their consumption of protein (Fig 7d) aligned closely with what would be expected under random consumption (Fig 7c).

**Figure 7.**
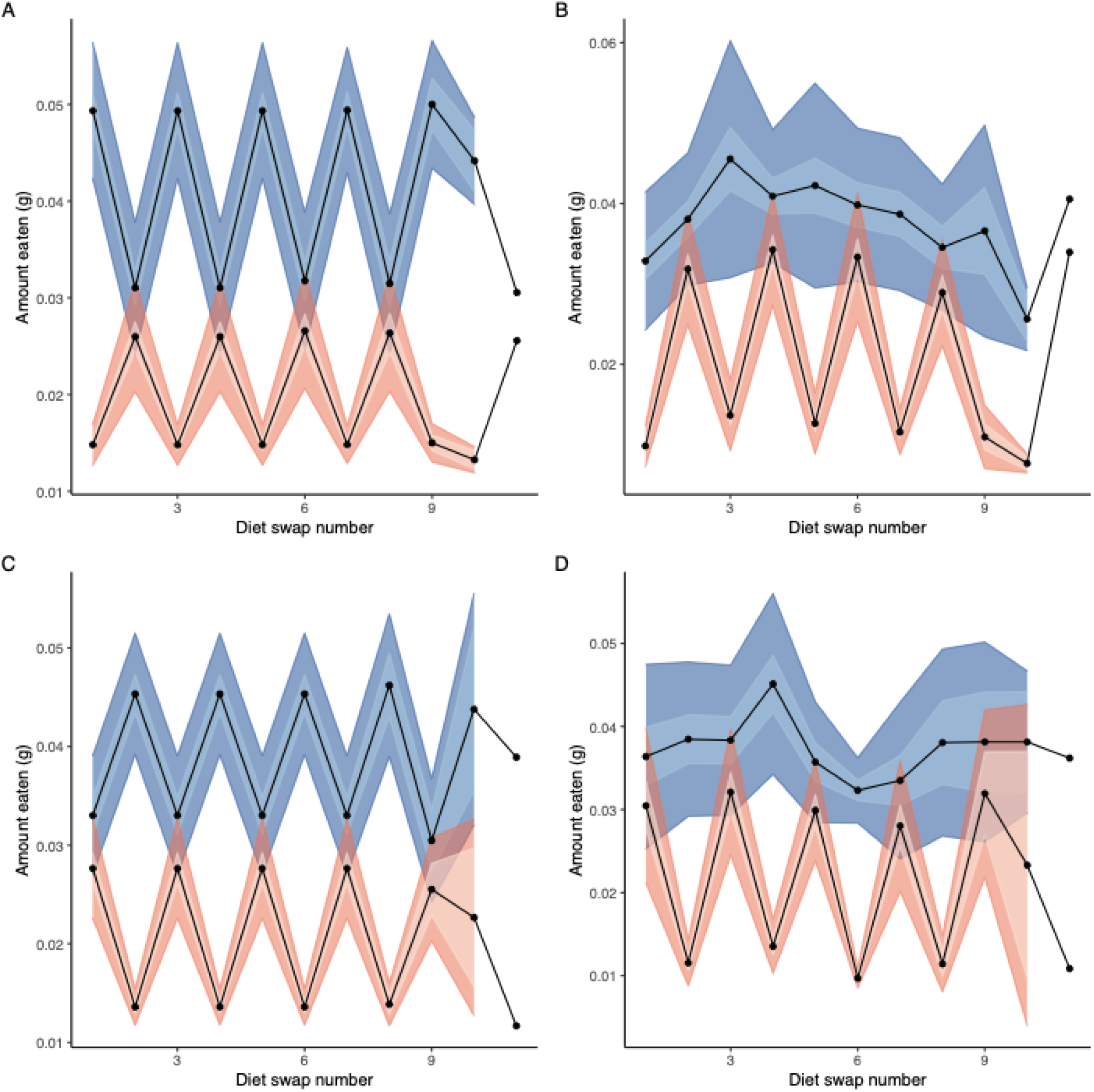
(a, c) Mean expected intake over successive diet swaps assuming random consumption of diets (+/-1 SE, inner ribbon, and SD, outer ribbon) of protein (red, lower ribbons) and carbohydrate (blue, upper ribbons), irrespective of the concentrations of the diet choices, for larvae starting on (a) diet A or (c) diet C. (b, d) Nutrient intake actually observed for larvae starting on (b) diet A or (d) diet C (+/-1 SE, inner ribbon, and SD, outer ribbon). For details of calculations of expected consumption, see text. Swap 11 lacks confidence intervals because only one larva in each group reached this stage.

## Discussion

We found that carbohydrate was positively associated with both body size and survival in *Osmia bicornis* larvae (Figs. 4a, 5a), although within our range of dietary ratios we did not specifically identify an optimum (fitness-maximising peak) in intake for either carbohydrate or protein. Accordingly, given a choice, larvae converged on a relatively carbohydrate-biased protein:carbohydrate ratio of 1:1.8 (Fig. 6a). Moreover, larvae prioritised carbohydrate over protein intake, showing tighter control over carbohydrate consumption than over protein consumption (Fig 7), and they pupated after eating about 0.23 g carbohydrate irrespective of protein and of dietary dilution (Fig 3). Yet this carbohydrate target fell short of the amount of carbohydrate that maximised cocoon weight or survival to pupation (Fig 4a, 5a). Dietary dilution imposed costs upon larvae regardless of nutritional intake, in the form of greater mortality and lower cocoon weights (Figs 4b, 5b). Taken together, these results show that (1) larval *O. bicornis* are at least partially responsible for their own nutritional regulation, and (2) their performance and consumption rules suggest adaptation to a pattern of carbohydrate-limited growth and survival. In what follows, we suggest how and why these patterns in *O. bicornis* depart from expected results based on studies of related organisms, and more generally what these findings suggest about nutritional cooperation and/or conflict between parents and offspring in (mass) provisioning species.

Larvae grew and survived best on our highest carbohydrate (i.e. lowest protein:carbohydrate) diets. Accordingly, across our range of diets, larvae maintained a constant carbohydrate intake while tolerating excesses or deficiencies of protein (a “no-interaction” rule of compromise; Raubenheimer & Simpson 1999b) - although it is conceivable that alternative rules of compromise, such as the “equal distance rule” more typically seen in generalist herbivores (Raubenheimer & Simpson 1999b; Behmer 2009), might have been evident over a broader array of diets. Both these patterns are unusual because insect herbivores are generally considered to be limited by protein (e.g. Bernays & Chapman 2007; although see Le Gall & Behmer 2014). In the few existing studies involving larval bees, e.g. honeybees (Helm *et al*. 2017), bumblebees (Kraus *et al*. 2019) and sweat bees (Roulston & Cane 2002), protein and not carbohydrate mediated larval growth and/or survival. More broadly, insect larvae often grow and survive best on balanced or moderately high protein:carbohydrate ratios (Roeder & Behmer 2014; Rodrigues *et al*. 2015), although low protein:carbohydrate ratios are associated with longevity in adults (e.g. Lee *et al*. 2008). Moreover, animals generally prioritise regulation of the nutrient that is typically limiting in their normal diet, and tolerate variation in nutrients that are abundant (Raubenheimer & Simpson 1999a). Tolerance of wide variation in protein is thus usually seen in predators (e.g. Raubenheimer *et al*. 2007; Kohl, Coogan & Raubenheimer 2015). In contrast, herbivores often regulate protein more tightly than carbohydrate (Lee *et al*. 2002; Le Gall & Behmer 2014; VanOverbeke, Thompson & Redak 2017). It is worth noting that protein *did* weakly mediate survival in our larvae to some extent, although only at low carbohydrate - possibly as a result of switching to protein as an energy source.

During the choice phase, when allowed to self-select diets, *O. bicornis* larvae converged on a protein:carbohydrate ratio of 1:1.8. This ratio is considerably more carbohydrate-biased than that preferred by bumblebees foraging on behalf of microcolonies (1:0.25, Vaudo *et al*. 2018; 1:0.08, Kraus *et al*. 2019), and (to a lesser extent) than ants foraging for colonies with offspring (1:1.5, Dussutour & Simpson 2009). It is also more carbohydrate-biased than that selected by reproductive, solitary phytophages such as grasshopper adults and lepidopteran larvae (1:0.25 - 1:1.4, reviewed in Behmer 2009) and is closer to diets selected by *Drosophila* larvae (1:2, Rodrigues *et al*. 2015). Notably, though, 1:1.8 was more protein-biased than the ratio that we found maximised both cocoon weight and survival (1:3.4), suggesting *O. bicornis* larvae may choose diets that favour other fitness-related quantities (such as reproduction and/or developmental time) over body size/survival, as in *Drosophila* (Lee *et al*. 2008; Rodrigues *et al*. 2015). As a cautionary note, the specific source of nutrients may also affect the preferred ratio: for example, adult honeybees exhibited different target P:C ratios when fed different protein sources (Altaye *et al*. 2010). Whether larvae are similarly sensitive is still unknown.

Two main features of *O. bicornis’* ecology may help to explain their prioritization of carbohydrate, and their relative preference for this macronutrient, compared to what we know of related taxa. First, the relative paucity of carbohydrate in *O. bicornis’* larval diet may help to explain these findings. Despite being herbivorous, *Osmia* larvae are unlikely to be protein-limited, because pollen is among the most protein-rich of plant tissues (Mattson 1980). Moreover, in *Osmia* specifically, nectar constitutes only a tiny fraction of the pollen ball, less than 4% (Maddocks & Paulus 1987; see Radmacher & Strohm 2010), in contrast to many other bees where nectar is a principal source of carbohydrate for larvae (e.g. Kraus *et al*. 2019). *O. bicornis* larvae may therefore be limited more by the amount of digestible carbohydrate within pollen than by dietary protein (see Roulston & Cane 2000). Second, *O. bicornis* is (to our knowledge) the first truly solitary hymenopteran studied under the GF; other studies have concerned individuals likely to become workers of social species. Unlike social hymenopterans, *O. bicornis* offspring are all reproductive and undergo diapause (Fliszkiewicz *et al*. 2012) - both activities dependent on the fat body, where carbohydrate-derived fat is stored (Kawooya & Law 1988; Ziegler & Van Antwerpen 2006; Hahn & Denlinger 2007; Wasielewski *et al*. 2013). Thus, *O. bicornis* larvae may have additional requirements for carbohydrate over and above those of developing nonreproductive, nondiapausing hymenopteran workers. These contrasting findings reinforce the idea that bees’ nutritional needs may be just as diverse as their ecologies.

Although larvae retained the ability to regulate carbohydrate by over- or under-eating protein, they nevertheless coped very poorly with dietary dilution (Fig 4b, 5b), despite displaying compensatory feeding behaviour (Fig 2) that suggests they both detected and responded to such dilution. The dilution was not excessive (70% nutrient density) compared to similar studies offering very highly dilute diets (14%, Raubenheimer & Simpson 1993; 16.8%, Lee, Raubenheimer & Simpson 2004). The locusts and caterpillars in those studies, though, are adapted for diets that vary greatly in nutrient density, beginning dilute and becoming even more dilute over the season (Scriber & Slansky 1981). By contrast, pollen is among the most consistently nutrient-rich parts of a plant (Roulston & Cane 2000) and does not broadly vary in composition over a season (DeGrandi-Hoffman *et al*. 2018). With a normal diet of unadulterated pollen and very little nectar, *Osmia* larvae may have had no need to evolve mechanisms to cope with dilution. In comparison, caterpillars reared on an invariant diet for generations lost the ability both to regulate intake and to cope with dilution (Warbrick-Smith *et al*. 2009). *Osmia* larvae appear to have retained the former capacity, but lost the latter, suggesting a normal diet that is dense in nutrients, but variable in composition.

In systems where parents gather food for offspring from the environment, both parents and offspring can be active participants in nutritional regulation. The lack of protein regulation shown by *O. bicornis* larvae highlights the importance of understanding (a) whether mother *O. bicornis* adjust protein content of provisions in response to imbalances in the landscape, and (b) whether larvae have physiological adaptations (e.g. post-ingestive processing) for tolerating protein imbalance. Budde & Lunau (2007) found that *O. bicornis* provisions contained about 19% protein regardless of pollen species used, suggesting a degree of homeostasis by parents. Yet human activity is reducing floral diversity and quality (Ziska *et al*. 2016; Papanikolaou *et al*. 2017). Evidence is mixed concerning whether, in practice, parent bees assess pollen nutrients at the flower (reviewed by Nicholls & Hempel de Ibarra 2016). Both bumblebees and ants balance nutrition on behalf of colonies (Dussutour & Simpson 2009; Vaudo *et al*. 2018), regulating more tightly when foraging for offspring - protein in the case of both taxa (Dussutour & Simpson 2009; Kraus *et al*. 2019) and carbohydrate in ants (Dussutour & Simpson 2008; Cook *et al*. 2010). On the other hand, protein gathered by honeybees varies passively with landscape usage (while maintaining carbohydrate and lipid; Donkersley *et al*. 2014). Which regulatory strategy *Osmia* parents and larvae collectively pursue may have important implications for their vulnerability to human-induced landscape change, and so should now be a focus for research.

Additionally, the ability to discriminate among nutrients provided by parents may be one tool offspring can use to exert some control over their nutrition, even in the absence of contact with parents. *Osmia* parents may provide suboptimal resources simply because of inefficiency in gathering pollen: efficiency drops across the season (Seidelmann 2006) and is lower in smaller-bodied parents (Seidelmann, Ulbrich & Mielenz 2010). Moreover, less efficient parents actively switch to producing male offspring (Seidelmann *et al*. 2010), so male and female offspring may experience different selection for regulation. This is well documented in other groups (e.g. Maklakov *et al*. 2008) and sex differences in larval regulation should now be a focus for research. But it is also well known that the evolutionary interests of parents and offspring frequently differ over how resources should be allocated (Trivers 1974; Crespi & Semeniuk 2004; Kilner & Drummond 2007; Haig 2010). The potential for offspring to use nutritional regulation to mitigate parentally imposed costs has been relatively overlooked, as most research to date has focused on parent-offspring conflict over *amount* of parental provisions, despite clear potential for conflict over composition (e.g. in discus fish, Buckley *et al*. 2010). Among primitively social Hymenoptera, some parents actively stunt offspring by restricting provisions (Lawson *et al*. 2017), securing their help by forcing them to become workers (Craig 1983). But the composition of food provided by parents is also critical to offspring fitness (e.g. Roulston & Cane 2002) and in extreme cases caste-determining (Anderson 1984). *O. bicornis* are solitary and lack castes, but this does not preclude parent-offspring disagreement over the optimal balance of offspring nutrition, as in e.g. *Drosophila* (Rodrigues *et al*. 2015).

More broadly, understanding the relative roles of offspring (intake regulation and post-ingestive processing) versus parents in nutrient balancing, as well as their evolutionary interests, will be key to understanding the nutritional ecology of species with parental provisioning. Such species include not just bees and other Hymenoptera, but other important ecosystem service providers such as dung beetles (Frank *et al*. 2017) and burying beetles (Hopwood, Moore & Royle 2013), as well as altricial birds (Wiens & Johnston 2012) and even humans (Burt & Amin 2014). Alloregulation by parents is not a given; the relative roles and interests of parent and offspring in these groups are likely to reflect species’ ecologies. Recent studies have found nutritional mismatches between oviposition sites selected by parents and the nutritional requirements of the offspring that will develop in those sites (Rodrigues *et al*. 2015; Lihoreau *et al*. 2016). Parent sweat bees (*Lasioglossum zephyrum*) appear not to regulate protein in larval provisions, despite protein mediating offspring performance (Roulston & Cane 2002). In *O. bicornis*, we have shown that offspring retain the ability to regulate their nutritional intake despite all food selection being done by parents whom they never meet. Larvae appeared to pay closest attention to regulating dietary carbohydrate, consistent with this nutrient mediating both growth and survival. Yet protein remains a key requirement for development; key now is to (a) establish the nutritional rules used by parents when provisioning offspring, and whether these coincide with or depart from those employed by larvae, and (b) establish specifically how protein balance is achieved, and whether parents or larvae carry that responsibility.

## Acknowledgements

The authors thank Victor Swetez, Emma Chapman, Toby Bagnall, Fiona Tainsh and Shannon Woodmansey for logistical assistance; Stephen Simpson, Audrey Dussutour, Mathieu Lihoreau, Michal Filipiak, Sheena Cotter, Lori Lawson Handley, Domino Joyce, Elizabeth Duncan, Francis Gilbert and Lucy Browning for valuable discussions and comments on the manuscript. The study was funded by the University of Hull; JDJG was also initially supported in this work by a research grant from the Association for the Study of Animal Behaviour.

## Author Contributions

JDJG and AA conceived and designed the study. AA gathered all data. Both authors analysed the data, and wrote and edited the manuscript.

## Data Accessibility

All data in this study will be made available upon acceptance via a digital repository such as DRYAD, Zenodo, Open Science Framework etc.

## References

Altaye, S.Z., Pirk, C.W.W., Crewe, R.M. & Nicolson, S.W. (2010) Convergence of carbohydrate-biased intake targets in caged worker honeybees fed different protein sources. The Journal of experimental biology, 213, 3311–3318.

Anderson, M. (1984) The evolution of eusociality. Annual review of ecology and systematics, 15, 165–189.

Behmer, S.T. (2009) Insect herbivore nutrient regulation. Annual review of entomology, 54, 165–187.

Bernays, E.A. & Chapman, R.F. (2007) Host-Plant Selection by Phytophagous Insects. Springer Science & Business Media.

Buckley, J., Maunder, R.J., Foey, A., Pearce, J., Val, A.L. & Sloman, K.A. (2010) Biparental mucus feeding: a unique example of parental care in an Amazonian cichlid. The Journal of experimental biology, 213, 3787–3795.

Budde, J. & Lunau, K. (2007) Rezepte für ein Pollenbrot-heute: Osmia rufa. Entomologie heute, 19, 173–179.

Burt, N.M. & Amin, M. (2014) A mini me?: exploring early childhood diet with stable isotope ratio analysis using primary teeth dentin. Archives of oral biology, 59, 1226–1232.

Cane, J.H. (2016) Adult Pollen Diet Essential for Egg Maturation by a Solitary Osmia Bee. Journal of insect physiology, 95, 105–109.

Cook, S.C., Eubanks, M.D., Gold, R.E. & Behmer, S.T. (2010) Colony-level macronutrient regulation in ants: mechanisms, hoarding and associated costs. Animal behaviour, 79, 429–437.

Costa, J.T. (2006) The Other Insect Societies. Harvard University Press.

Craig, R. (1983) Subfertility and the evolution of eusociality by kin selection. Journal of theoretical biology, 100, 379–397.

Crespi, B. & Semeniuk, C. (2004) Parent-offspring conflict in the evolution of vertebrate reproductive mode. The American naturalist, 163, 635–653.

Deans, C., Sword, G.A. & Behmer, S.T. (2019) First evidence of protein-carbohydrate regulation in a plant bug (Lygus hesperus). Journal of insect physiology, 116, 118–124.

DeGrandi-Hoffman, G., Gage, S.L., Corby-Harris, V., Carroll, M., Chambers, M., Graham, H., Watkins deJong, E., Hidalgo, G., Calle, S., Azzouz-Olden, F., Meador, C., Snyder, L. & Ziolkowski, N. (2018) Connecting the nutrient composition of seasonal pollens with changing nutritional needs of honey bee (Apis mellifera L.) colonies. Journal of insect physiology, 109, 114–124.

Despland, E. & Noseworthy, M. (2006) How well do specialist feeders regulate nutrient intake? Evidence from a gregarious tree-feeding caterpillar. The Journal of experimental biology, 209, 1301–1309.

Donkersley, P., Rhodes, G., Pickup, R.W., Jones, K.C. & Wilson, K. (2014) Honeybee nutrition is linked to landscape composition. Ecology and evolution, 4, 4195–4206.

Dussutour, A. & Simpson, S.J. (2008) Carbohydrate regulation in relation to colony growth in ants. The Journal of experimental biology, 211, 2224–2232.

Dussutour, A. & Simpson, S.J. (2009) Communal nutrition in ants. Current biology: CB, 19, 740–744.

Falk, S.J. (2015) Field Guide to the Bees of Great Britain and Ireland. British Wildlife Publishing.

Fichter, B.L., Stephen, W.P. & Vandenberg, J.D. (1981) An Aseptic Technique for Rearing Larvae of the Leafcutting Bee Megachile Rotundata (Hymenoptera, Megachilidae). Journal of apicultural research, 20, 184–188.

Field, J. (2005) The evolution of progressive provisioning. Behavioral ecology, 16 (4), 770–778

Filipiak, M. (2019) Key pollen host plants provide balanced diets for wild bee larvae: A lesson for planting flower strips and hedgerows (ed R Rader). The Journal of applied ecology, 56 (6), 1410–1418

Fliszkiewicz, M., Giejdasz, K., Wasielewski, O. & Krishnan, N. (2012) Influence of winter temperature and simulated climate change on body mass and fat body depletion during diapause in adults of the solitary bee, Osmia rufa (Hymenoptera: Megachilidae). Environmental entomology, 41, 1621–1630.

Frank, K., Brückner, A., Hilpert, A., Heethoff, M. & Blüthgen, N. (2017) Nutrient quality of vertebrate dung as a diet for dung beetles. Scientific reports, 7, 12141.

Goulson, D., Nicholls, E., Botías, C. & Rotheray, E.L. (2015) Bee declines driven by combined stress from parasites, pesticides, and lack of flowers. Science, 347,1255957.

Hahn, D.A. & Denlinger, D.L. (2007) Meeting the energetic demands of insect diapause: nutrient storage and utilization. Journal of insect physiology, 53, 760–773.

Haider, M., Dorn, S., Sedivy, C. & Muller, A. (2014) Phylogeny and floral hosts of a predominantly pollen generalist group of mason bees (Megachilidae: Osmiini). Biological journal of the Linnean Society. Linnean Society of London, 111, 78–91.

Haig, D. (2010) Colloquium papers: Transfers and transitions: parent-offspring conflict, genomic imprinting, and the evolution of human life history. Proceedings of the National Academy of Sciences of the United States of America, 107 Suppl 1, 1731–1735.

Harper, E.J. & Turner, C.L. (2000) Nutrition and energetics of the canary (Serinus canarius). Comparative biochemistry and physiology. Part B, Biochemistry & molecular biology, 126, 271–281.

Helm, B.R., Slater, G.P., Rajamohan, A., Yocum, G.D., Greenlee, K.J. & Bowsher, J.H. (2017) The geometric framework for nutrition reveals interactions between protein and carbohydrate during larval growth in honey bees. Biology open, 6, 872–880.

Hogendoorn, K. & Velthuis, H.H.W. (1993) The sociality of Xylocopa pubescens: does a helper really help? Behavioral ecology and sociobiology, 32, 247–257.

Hopwood, P.E., Moore, A.J. & Royle, N.J. (2013) Nutrition during sexual maturation affects competitive ability but not reproductive productivity in burying beetles (ed W Blanckenhorn). Functional ecology, 27, 1350–1357.

Hunt, J.H. & Nalepa, C.A. (1994) Nourishment and Evolution in Insect Societies. Westview Press.

Jauker, F., Bondarenko, B., Becker, H.C. & Steffan-Dewenter, I. (2012) Pollination efficiency of wild bees and hoverflies provided to oilseed rape. Agricultural and forest entomology, 14, 81–87.

Kawooya, J.K. & Law, J.H. (1988) Role of lipophorin in lipid transport to the insect egg. The Journal of biological chemistry, 263, 8748–8753.

Kilner, R.M. & Drummond, H. (2007) Parent--offspring conflict in avian families. Journal of ornithology, 148, 241–246.

Kohl, K.D., Coogan, S.C.P. & Raubenheimer, D. (2015) Do wild carnivores forage for prey or for nutrients? BioEssays: news and reviews in molecular, cellular and developmental biology, 37, 701–709.

Kraus, S., Gómez-Moracho, T., Pasquaretta, C., Latil, G., Dussutour, A. & Lihoreau, M. (2019) Bumblebees adjust protein and lipid collection rules to the presence of brood. Current zoology, 65, 437–446.

Lawson, S.P., Helmreich, S.L. & Rehan, S.M. (2017) Effects of nutritional deprivation on development and behavior in the subsocial bee Ceratina calcarata (Hymenoptera: Xylocopinae). The Journal of experimental biology, 220, 4456–4462.

Lee, K.P., Behmer, S.T., Simpson, S.J. & Raubenheimer, D. (2002) A geometric analysis of nutrient regulation in the generalist caterpillar Spodoptera littoralis (Boisduval). Journal of insect physiology, 48, 655–665.

Lee, K.P., Raubenheimer, D. & Simpson, S.J. (2004) The effects of nutritional imbalance on compensatory feeding for cellulose-mediated dietary dilution in a generalist caterpillar. Physiological entomology, 29, 108–117.

Lee, K.P., Simpson, S.J., Clissold, F.J., Brooks, R., Ballard, J.W.O., Taylor, P.W., Soran, N. & Raubenheimer, D. (2008) Lifespan and reproduction in Drosophila: New insights from nutritional geometry. Proceedings of the National Academy of Sciences of the United States of America, 105, 2498–2503.

Le Gall, M. & Behmer, S.T. (2014) Effects of protein and carbohydrate on an insect herbivore: the vista from a fitness landscape. Integrative and comparative biology, 54, 942–954.

Lihoreau, M., Buhl, J., Charleston, M.A., Sword, G.A., Raubenheimer, D. & Simpson, S.J. (2014) Modelling nutrition across organizational levels: from individuals to superorganisms. Journal of insect physiology, 69, 2–11.

Lihoreau, M., Poissonnier, L.-A., Isabel, G. & Dussutour, A. (2016) Drosophila females trade off good nutrition with high-quality oviposition sites when choosing foods. The Journal of experimental biology, 219, 2514–2524.

Mackenzie, G., Boa, A.N., Diego-Taboada, A., Atkin, S.L. & Sathyapalan, T. (2015) Sporopollenin, The Least Known Yet Toughest Natural Biopolymer. Frontiers of materials science, 2, 129.

Maddocks, R. & Paulus, H.F. (1987) Quantitative Aspekte der Brut-biologie von Osmia rufa L. und Osmia cornuta Latr.(Hymenoptera, Megachilidae): Eine vergleichende Untersuchung zu Mechanismen der Konkurrenzminderunt zweier nahverwandter Bienenarten. Zoologische Jahrbücher. Abteilung für Systematik, Ökologie und Geographie der Tiere, 114, 15–44.

Maklakov, A.A., Simpson, S.J., Zajitschek, F., Hall, M.D., Dessmann, J., Clissold, F., Raubenheimer, D., Bonduriansky, R. & Brooks, R.C. (2008) Sex-specific fitness effects of nutrient intake on reproduction and lifespan. Current biology, 18, 1062–1066.

Mattson, W.J. (1980) Herbivory in Relation to Plant Nitrogen Content. Annual review of ecology and systematics, 11, 119–161.

Mayntz, D., Raubenheimer, D., Salomon, M., Toft, S. & Simpson, S.J. (2005) Nutrient-specific foraging in invertebrate predators. Science, 307, 111–113.

Michaelsen, K.F., Weaver, L, Branca, F. & Robertson, A. (2003) Feeding and Nutrition of Infants and Young Children. WHO Regional Publications, European Series.

Nelson, E.V., Roberts, R.B. & Stephen, W.P. (1972) Rearing Larvae of the Leaf-Cutter Bee Megachile Rotundata on Artificial Diets. Journal of apicultural research, 11, 153–156.

Nicholls, E. & Hempel de Ibarra, N. (2016) Assessment of pollen rewards by foraging bees. Functional ecology, 31, 76–87.

Papanikolaou, A.D., Kühn, L, Frenzel, M., Kuhlmann, M., Poschlod, P., Potts, S.G., Roberts, S.P.M. & Schweiger, O. (2017) Wild bee and floral diversity co-vary in response to the direct and indirect impacts of land use. Ecosphere, 8 (11), e02008.

Poissonnier, L.-A., Arganda, S., Simpson, S.J., Dussutour, A. & Buhl, J. (2018) Nutrition in extreme food specialists: An illustration using termites. Functional ecology, 32, 2531–2541.

Radmacher, S. & Strohm, E. (2010) Factors affecting offspring body size in the solitary bee Osmia bicornis (Hymenoptera, Megachilidae). Apidologie, 41, 169–177.

Raubenheimer, D. & Jones, S.A. (2006) Nutritional imbalance in an extreme generalist omnivore: tolerance and recovery through complementary food selection. Animal behaviour, 71, 1253–1262.

Raubenheimer, D., Mayntz, D., Simpson, S.J. & Tøft, S. (2007) Nutrient-specific compensation following diapause in a predator: implications for intraguild predation. Ecology, 88 (10), 2598–2608.

Raubenheimer, D. & Simpson, S.J. (1993) The geometry of compensatory feeding in the locust. Animal behaviour, 45, 953–964.

Raubenheimer, D. & Simpson, S.J. (1999a) Integrating nutrition: a geometrical approach. Proceedings of the 10th International Symposium on Insect-Plant Relationships, Series Entomologica, pp. 67–82. Springer Netherlands.

Raubenheimer, D. & Simpson, S.J. (1999b) Integrating nutrition: a geometrical approach. Entomologia experimentalis et applicata, 91, 67–82.

Raubenheimer, D., Simpson, S.J. & Mayntz, D. (2009) Nutrition, ecology and nutritional ecology: toward an integrated framework. Functional ecology, 23, 4–16.

Rehan, S.M., Richards, M.H., Adams, M. & Schwarz, M.P. (2014) The costs and benefits of sociality in a facultatively social bee. Animal behaviour, 97, 77–85.

Rodrigues, M.A., Martins, N.E., Balancé, L.F., Broom, L.N., Dias, A.J.S., Fernandes, A.S.D., Rodrigues, F., Sucena, É. & Mirth, C.K. (2015) Drosophila melanogaster larvae make nutritional choices that minimize developmental time. Journal of insect physiology, 81, 69–80.

Roeder, K.A. & Behmer, S.T. (2014) Lifetime consequences of food protein-carbohydrate content for an insect herbivore (ed G Davidowitz). Functional ecology, 28, 1135–1143.

Roulston, T.H. & Cane, J.H. (2000) Pollen nutritional content and digestibility for animals. Plant systematics and evolution = Entwicklungsgeschichte und Systematik der Pflanzen, 222, 187–209.

Roulston, T.H. & Cane, J.H. (2002) The effect of pollen protein concentration on body size in the sweat bee Lasioglossum zephyrum (Hymenoptera: Apiformes). Evolutionary ecology, 16, 49–65.

Roulston, T.H. & Goodell, K. (2011) The role of resources and risks in regulating wild bee populations. Annual review of entomology, 56, 293–312.

Royama, T. (1970) Factors governing the hunting behaviour and selection of food by the great tit (Parus major. The Journal of animal ecology, 39, 619–668.

Schmickl, T. & Karsai, I. (2017) Resilience of honeybee colonies via common stomach: A model of self-regulation of foraging. PloS one, 12, e0188004.

Scriber, J.M. & Slansky, F., Jr. (1981) The Nutritional Ecology of Immature Insects. Annual review of entomology, 26, 183–211.

Seidelmann, K. (2006) Open-cell parasitism shapes maternal investment patterns in the Red Mason bee Osmia rufa. Behavioral ecology: official journal of the International Society for Behavioral Ecology, 17, 839–848.

Seidelmann, K., Ulbrich, K. & Mielenz, N. (2010) Conditional sex allocation in the Red Mason bee, Osmia rufa. Behavioral ecology and sociobiology, 64, 337–347.

Simpson, S.J. & Raubenheimer, D. (1993) A Multi-Level Analysis of Feeding Behaviour: The Geometry of Nutritional Decisions. Philosophical transactions of the Royal Society of London. Series B, Biological sciences, 342, 381–402.

Simpson, S.J. & Raubenheimer, D. (2012) The nature of nutrition: a unifying framework. Australian journal of zoology, 59, 350–368.

Smiseth, P.T., Wright, J. & Kölliker, M. (2008) Parent-offspring conflict and co-adaptation: behavioural ecology meets quantitative genetics. Proceedings. Biological sciences / The Royal Society, 275, 1823–1830.

Strohm, E., Daniels, H., Warmers, C. & Stoll, C. (2002) Nest provisioning and a possible cost of reproduction in the megachilid bee Osmia rufa studied by a new observation method. Ethology Ecology & Evolution, 14, 255–268.

Tainsh, F., Woodmansey, S.R., Austin, A.J., Bagnall, T.E. & Gilbert, J.D.J. (2020) Sporopollenin as a dilution agent in artificial diets for solitary bees. Apidologie, https://doi.org/10.1007/s13592-020-00801-1

Trivers, R.L. (1974) Parent-Offspring Conflict. Integrative and comparative biology, 14, 249–264.

VanOverbeke, D.R., Thompson, S.N. & Redak, R.A. (2017) Dietary self-selection and rules of compromise by fifth-instar Vanessa cardui. Entomologia experimentalis et applicata, 163, 209–219.

Vaudo, A.D., Farrell, L.M., Patch, H.M., Grozinger, C.M. & Tooker, J.F. (2018) Consistent pollen nutritional intake drives bumble bee (Bombus impatiens) colony growth and reproduction across different habitats. Ecology and evolution, 8 (11), 5765–5776

Vaudo, A.D., Patch, H.M., Mortensen, D.A., Tooker, J.F. & Grozinger, C.M. (2016) Macronutrient ratios in pollen shape bumble bee (Bombus impatiens) foraging strategies and floral preferences. Proceedings of the National Academy of Sciences of the United States of America, 113, E4035–42.

Warbrick-Smith, J., Raubenheimer, D., Simpson, S.J. & Behmer, S.T. (2009) Three hundred and fifty generations of extreme food specialisation: testing predictions of nutritional ecology. Entomologia experimentalis et applicata, 132, 65–75.

Wasielewski, O., Wojciechowicz, T., Giejdasz, K. & Krishnan, N. (2013) Overwintering strategies in the red mason solitary bee—physiological correlates of midgut metabolic activity and turnover of nutrient reserves in females of Osmia bicornis. Apidologie, 44, 642–656.

Weeks, R.D., Wilson, L.T., Vinson, S.B. & James, W.D. (2004) Flow of Carbohydrates, Lipids, and Protein Among Colonies of Polygyne Red Imported Fire Ants, Solenopsis invicta (Hymenoptera: Formicidae). Annals of the Entomological Society of America, 97, 105–110.

Wiens, J. & Johnston, R. (2012) Adaptive correlates of granivory in birds. Granivorous Birds in Ecosystems: Their Evolution, Populations, Energetics, Adaptations, Impact and Control,. (Eds J. Pinowski and SC Kendeigh.) pp, 301–340.

Ziegler, R. & Van Antwerpen, R. (2006) Lipid uptake by insect oocytes. Insect biochemistry and molecular biology, 36, 264–272.

Ziska, L.H., Pettis, J.S., Edwards, J., Hancock, J.E., Tomecek, M.B., Clark, A., Dukes, J.S., Loladze, I. & Polley, H.W. (2016) Rising atmospheric CO2 is reducing the protein concentration of a floral pollen source essential for North American bees. Proceedings. Biological sciences / The Royal Society, 283 (1828), 20160414

