## Supplementary material for "Solitary bee larvae prioritize carbohydrate over protein in parentally provided pollen": sporopollenin prep, nesting materials, larval removal (from another version of this preprint: version 2, 2020-08-18)

*S1. Pollen husk (sporopollenin) preparation*

To produce the exines 500g of raw sunflower pollen (Henan Shengchoa Apiculture Co., China) was suspended in 9M aqueous HCL solution (2250ml) and heated at 94-97°C for one hour. The solution was then neutralised with a 50% KOH solution, the resulting fluid filtered out using a Buchner funnel under vacuum, and the filtrate discarded. The residue was then washed with water, methanol (100%), and acetone (100%) sequentially under vacuum to remove any remaining solution. Each wash was repeated until the effluent ran clear before moving to the next washing stage i.e. filtrate was first washed with water until effluent ran clear, then methanol etc. Once the final wash was completed, filtrate was allowed to dry overnight. These 'husks' were then subjected to SEM imaging to ensure that only the outer pollen shell, the husk, remained.

*S2. Nesting materials and larval removal*

The nesting material used to raise *O.bicornis* larvae consisted of Styrofoam blocks (Styrodur 3035 CS), with a 9x9mm furrow and polycarbonate lid, housed within a wooden frame (modified from Strohm et al., 2002; Fig. S1a,b). Completed nests, signified by a mud plug at the entrance, were then brought into the laboratory. Newly emerged larvae were left alone for two days before we transferred them to a single-occupancy nest and assigned each to an experimental treatment. The majority of nests were completed and sealed by the adult before the larvae hatched. When this was not the case, and some eggs hatched whilst the adult was still provisioning other cells in the nest, larvae reaching the two-day-post-hatching stage were removed at the nest site, taken to the lab and placed into a single-occupancy nest.

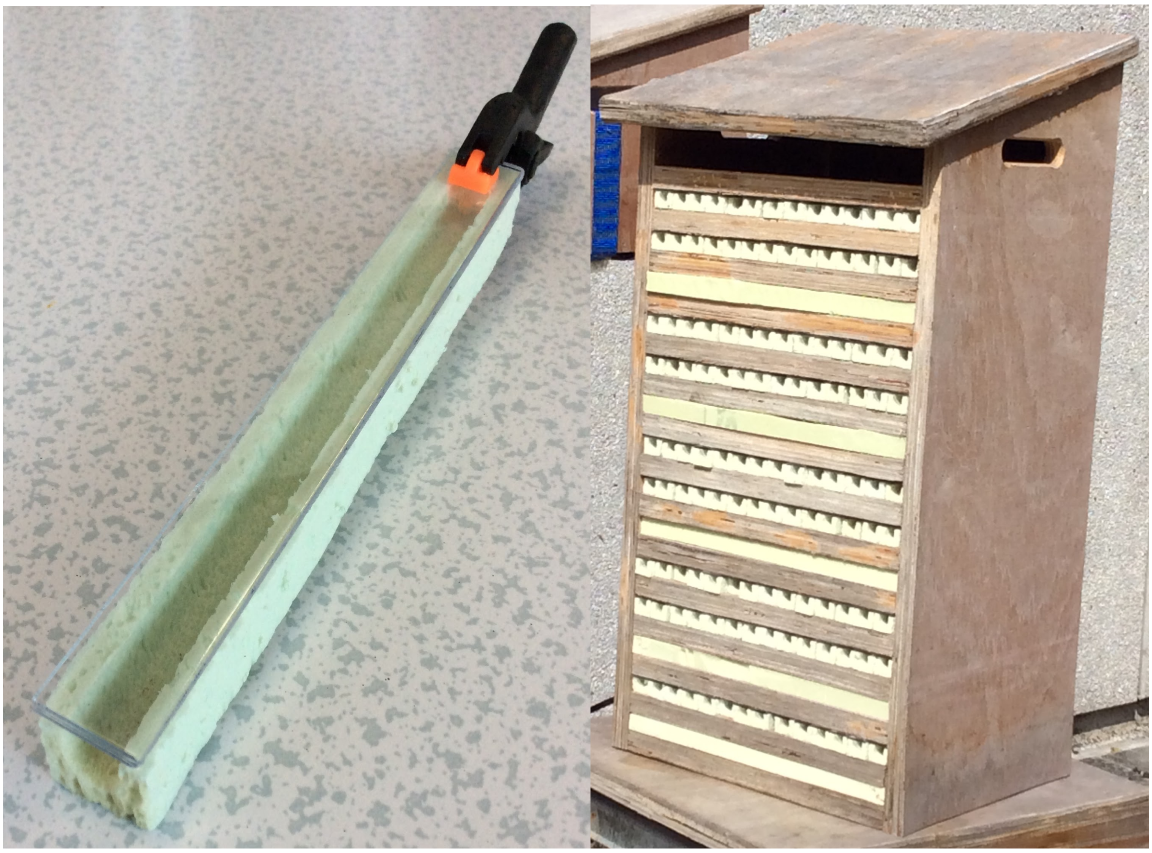

**Fig. S1. Nesting material provided to *O. bicornis* adults during 2017 season. A) Styrofoam (Styrodur, 3035CS) nest block with polycarbonate lid and clamp. B) Wooden housing containing nesting material.**
